# Global Spread of SARS-CoV-2 Subtype with Spike Protein Mutation D614G is Shaped by Human Genomic Variations that Regulate Expression of *TMPRSS2* and *MX1* Genes

**DOI:** 10.1101/2020.05.04.075911

**Authors:** Chandrika Bhattacharyya, Chitrarpita Das, Arnab Ghosh, Animesh K. Singh, Souvik Mukherjee, Partha P. Majumder, Analabha Basu, Nidhan K. Biswas

## Abstract

COVID-19 pandemic is a major human tragedy. Worldwide, SARS-CoV-2 has already infected over 3 million and has killed about 230,000 people. SARS-CoV-2 originated in China and, within three months, has evolved to an additional 10 subtypes. One particular subtype with a non-silent (Aspartate to Glycine) mutation at 614^th^ position of the Spike protein (D614G) rapidly outcompeted other pre-existing subtypes, including the ancestral. We assessed that D614G mutation generates an additional serine protease (Elastase) cleavage site near the S1-S2 junction of the Spike protein. We also identified that a single nucleotide deletion (delC) at a known variant site (rs35074065) in a cis-eQTL of *TMPRSS2*, is extremely rare in East Asians but is common in Europeans and North Americans. The delC allele facilitates entry of the 614G subtype into host cells, thus accelerating the spread of 614G subtype in Europe and North America where the delC allele is common. The delC allele at the cis-eQTL locus rs35074065 of *TMPRSS2* leads to overexpression of both *TMPRSS2* and a nearby gene *MX1*. The cis-eQTL site, rs35074065 overlaps with a transcription factor binding site of an activator (*IRF1*) and a repressor (IRF2). IRF1 activator can bind to variant delC allele, but IRF2 repressor fails to bind. Thus, in an individual carrying the delC allele, there is only activation, but no repression. On viral entry, *IRF1* mediated upregulation of *MX1* leads to neutrophil infiltration and processing of 614G mutated Spike protein by neutrophil Elastase. The simultaneous processing of 614G spike protein by TMPRSS2 and Elastase serine proteases facilitates the entry of the 614G subtype into host cells. Thus, SARS-CoV-2, particularly the 614G subtype, has spread more easily and with higher frequency to Europe and North America where the delC allele regulating expression of *TMPRSS2* and *MX1* host proteins is common, but not to East Asia where this allele is rare.

## Introduction

Coronaviruses infect humans with varying degrees of severity and lethality (*1*). Four of these viruses (NL63, 229E, OC43, and HKU1) cause mild respiratory problems in humans and three others (MERS-CoV, SARS-CoV including the newly emerged SARS-CoV-2) can cause severe respiratory syndromes (*2–4*). SARS-CoV-2 infection was first reported from Wuhan, China, on 24^th^ December 2019 (*2*) and in less than three months, on 11^th^ March 2020, WHO declared COVID-19 as a pandemic (*5*). By 3^rd^ May, 2020, 3.3 million people worldwide (213 countries) were reportedly infected and 2,38,628 individuals died of COVID-19 (*6*). We note that while the infectivity of SARS-CoV-2 is much higher than SARS-CoV or MERS-CoV (*7*), its case fatality rate (0.9-3.3%) is substantially lower than that of SARS-CoV (11%) and MERS-CoV (34%) (*8*). For SARS-CoV-2, there are notable differences in case-fatality rate and disease severity among geographical regions and among age groups of infected persons, with lower severity in infants and children than in adults (*1, 9*). The case fatality rates in East Asia (*10*) (China and neighbourhood) and Middle East (*11*) (Iran and neighbourhood) have been substantially lower than in Europe(*12*) (Italy, France and Spain).

Based on studies of its RNA sequence, it is clear that SARS-CoV-2 has evolved and diversified as it spread geographically (*13, 14*). RNA viruses acquire mutations easily. Most mutations are deleterious and viruses with mutations are eliminated. If a mutation attains a high frequency, it is expected that the mutation provides selective advantage to the virus, usually manifested by higher transmission efficiency. Viral RNA of SARS-CoV-2, collected and sequenced from isolates in different countries has resulted in the recognition of several subtypes (*15*). The ancestral type (O), likely to have originated in Wuhan, was first reported from China in late December 2019. Four months later, another type (A2a), which was also first reported from China on 24^th^ January 2020, spread rapidly and widely across Europe and North America [GISAID: https://www.gisaid.org/ and Nextstrain: https://nextstrain.org/] outcompeting the ancestral O type (*13, 16*).

SARS-CoV-2 virus is a single stranded (+) sense RNA virus with a genome length of about 30 Kb. It encodes 29 structural and non-structural proteins, including ORF1a/b polyprotein, spike (S) glycoprotein, envelope (E), membrane (M) and the nucleocapsid (N) protein (*8, 17*). To enable replication and spread, the virus binds to host cell surface receptors using spike protein to mediate fusion of the viral envelope with cell membrane. The spike glycoprotein consists of three S1-S2 heterodimers and it is present on the surface of the virus. The receptor binding domain (RBD) located on the head of the S1 domain of the viral Spike (S) protein attaches with the angiotensin converting enzyme 2 (ACE2), that is expressed in large quantities in specific cell types (pneumocytes) of the human lung, and other tissues (*18–20*). As long as the S1 and S2 subunits are parts of a single polypeptide chain, the S protein is inactive. Recent studies have shown that the type II transmembrane serine protease (TMPRSS2) of the host cleaves the Spike protein at the S1-S2 junction, where a furin cleavage site recognized by proteases is present, that facilitates entry of SARS-CoV-2 by membrane fusion involving the S2 domain (*21–24*). Functional studies on SARS-CoV showed that specific amino acid residues on the Spike protein are important determinants for virus tropism and transmission efficiency (*25*).

Variations in nucleotide sequences of *ACE2* and *TMPRSS2*, the two host genes whose products are indispensable (*21, 24, 26–28*) for the entry of the coronavirus into host cells, can alter the expression and functionality of these proteins. Recent analysis of single cells that express both ACE2 and TMPRSS2 has identified a ‘gene expression program’ in nasal, lung and other tissues that likely facilitates viral entry modulating interferon regulation assisted by host proteases (*18*). In *ACE2*, the majority of observed variants occur at low frequencies (non-polymorphic) in most human populations. Even though some recent studies (*29, 30*) have attempted to implicate these variants with susceptibility to infection by SARS-CoV-2, there is no convincing evidence yet that the low-frequency, non-polymorphic variants in *ACE2* can modulate susceptibility to SARS-Cov-2 infection. However, *TMPRSS2*, the product of which is involved in the proteolytic cleavage of both *ACE2* and spike proteins of SARS-CoV-2 leading to internalization of the virion in the host cell, harbors many variants that exhibit considerable variation in frequencies among human populations. We note that the geographical spread and increase in frequency of the A2a subtype (with a characteristic mutation, D614G, in the Spike protein) has been explosive (Nextstrain: https://nextstrain.org/). Compared to other clades, the frequency of A2a in Europe and North America, but not in East Asia, has increased rapidly to high levels (*13*). We surmise that host genomics may play a role in shaping the trajectory of rise and eventual level of the frequency of A2a in populations. The role of sequence variations in the *TMPRSS2* gene and other genomic regions that regulate the product of *TMPRSS2* has not been explored in the context of subtype specific transmission patterns of SARS-CoV-2. We hypothesize that gene variants that modulate expression of the serine protease TMPRSS2 in the human lung (non-silent variants of *TMPRSS2* and variants in its regulatory expression quantitative trait loci [eQTLs]), which is the major mediator of cellular entry of the SARS-CoV-2 virus, influence the infectivity of SARS-CoV-2 subtypes.

## Materials and Methods

### Phylodynamic Analysis of SARS-CoV-2 RNA sequences

One cardinal feature of the COVID-19 pandemic is our ability to monitor the spread and evolution of the SARS-CoV-2 virus almost in real time, since RNA sequences. of the virus are being deposited every day in large numbers from all global regions to public databases. In partial fulfilment of the objective of this study, we have analysed the most recent publicly available data. We downloaded all SARS-CoV-2 RNA sequences (n=8847) excluding low coverage (>5% N in the 29.9 Kb of each RNA sequence) on 16^th^ April 2020, 8:50 AM, from the GISAID database (*31*).

To analyze SARS-CoV-2 sequence data, we used the community standard nextstrain/ncov (*15*) (github.com/nextstrain/ncov) pipeline developed specifically for spatial and temporal tracking of pathogens. Nextstrain/ncov (*15*), is an open-source pipeline for phylodynamic analysis (*32*), including subsampling, alignment, phylogenetic inference, temporal dating of ancestral nodes and discrete trait geographic reconstruction as well as interactive data visualization. It comprises augur (*15*) (github.com/nextstrain/augur) a modular bioinformatics tool used for data analyses, and auspice (*15*) (github.com/nextstrain/auspice) a web-based visualization tool for phylogenomic and phylogeographic data.

Each downloaded fasta file was preprocessed to remove duplicate samples based on the identifier from the fasta headers (hCoV-19/<Country>/<Identifier>/<Year>). Out of the 8847 sequences, 6424 sequences passed default QC criteria of Nextstrain pipeline. These 6424 sequences were aligned using MAFFT (*33*). We estimated timescale and branch lengths of a reconstructed phylogenetic tree using IQ-TREE (*34*) (as implemented in augur) considering hCoV-19/Wuhan/WH01/2019 as ancestral (*31*); (https://www.gisaid.org/). These estimates were further refined using RAxML. Augur also estimates the frequency-trajectories of mutations, genotypes and clades of a phylogenetic tree, which is used by the auspice package to visually represent phylogenetic tree, geographic transmission and entropy (genetic diversity).

We used the date of viral sample collection for all phylodynamic analysis to understand epidemiological and evolutionary patterns. The initial sampling for sequencing was sparse, possibly non-representative and unstable. Therefore, to focus on a more stable virus infection period, we analyzed the data on 6181 out of 6424 viral RNA sequences that were collected between January 15, and March 31, 2020. Details of the phylogenetic tree of the viral sequences and defining mutations of various clades are provided in Supplementary Table 1. The isolate specific mutations, which occurred on the top branch of the tree, were identified from the results obtained by analyzing data on the Nextstrain pipeline. Temporal acquisition of subtype-specific mutations was also ascertained. Estimation of the number of segregating sites and values of Tajima’s D (*35*) for various clades were obtained using MEGA (*36*) and cross-validated with DNASP (*37*).

### In-Silico prediction of cleavage site on SARS-CoV-2 Spike protein

Functional studies showed that by synthetically introducing mutations near the S1-S2 junction of SARS-CoV additional proteolytic cleavage sites are generated that enhance viral membrane fusion by several fold (*38*). We downloaded the sequence of SARS-Cov-2 spike (S) protein (QHD43416), which is 1273 amino acids long, from https://zhanglab.ccmb.med.umich.edu/COVID-19/ (*39*). We used PROSPER (*40*) (https://prosper.erc.monash.edu.au/) to predict proteolytic cleavage sites based on i) local amino acid sequence profile, ii) predicted secondary structure, iii) solvent accessibility and iv) predicted native disorder. In particular we identified potential protease substrate sites in the amino acid sequence of SARS-CoV-2 spike (S) protein. The predicted protease cleavage sites were also verified by another protease cut site prediction tool PROSPERous (*41*).

### Tissue specific expression and genomic variation in human *TMPRSS2* in global populations

Using GTEx data (*42*) (https://gtexportal.org/), we identified regulatory eQTLs that are significantly associated with *TMPRSS2* gene expression in various human tissues. Genotype data, for all significant eQTLs as well as the data on all variants on the *TMPRSS2* gene, were extracted from 1000 Genomes dataset [https://ftp-trace.ncbi.nih.gov/1000genomes/ftp/release/20110521/] using tabix (*43*) with the hg19 chromosomal coordinates as reference. Initial data to assess population genomic diversity were downloaded from 1000 Genomes project that included representative populations from Europe (CEU, TSI, FIN, GBR, IBS), admixed Hispanic speakers from America (MXL, PUR, CLM) and East Asia (CHB, CHD, JPT) [Supplementary Table 2]. Functional annotation of each identified variant was done using Annovar (*44*). Apart from the eQTLs, only non-silent polymorphic variants in *TMPRSS2* that are more likely to cause alteration of protein structure and function, were selected for further analysis. We ranked the eQTLs according to their impact on the expression level of *TMPRSS2* in lung and selected variants with high impact. For all these eQTLs and non-silent variants, we calculated the genetic distance, F_st_ (*45*), between pairs of major continental populations and regional subpopulations, using PLINK v1.9 (*46*) (https://www.cog-genomics.org/plink/). Additional data on allele frequencies of non-silent variants in *TMPRSS2* gene and the highest ranked eQTL of *TMPRSS2* gene from 58 extant global populations, with n>10 individuals, were downloaded from multiple databases after careful data curation. The names of the population groups, geographic regions and database sources are provided in Supplementary Table 2.

### Transcription factor binding (TFB) prediction for human *TMPRSS2* eQTL sites

From the hg19 reference sequence, 20 nucleotides on each side of each eQTL variant site was considered for TFB prediction. At each site, for the two sequences containing the reference and the variant alleles, we used JASPAR (*47*) (http://jaspar.genereg.net/) to predict TF recognition sites with the default (80%) relative profile threshold.

## Results

### Temporal and geographical spread of subtypes of SARS-CoV-2

The first set of RNA sequences collected from 17 infected individuals from Wuhan, China, had high sequence identity, and was named the O subtype. The O subtype spread to other provinces (e.g., Guangdong, Jiangxi) of China and also to nearby countries, e.g., Thailand (with the first submission to GISAID – Nonthaburi/61/2020 – on 8^th^ January 2020). within two weeks. Our phylodynamic analysis showed that the virus evolved into B and B2 in the first two weeks of January and later to B1, B4, A2a and A3 [Supplementary Figure 1(a)]. These subtypes rapidly spread worldwide to multiple East Asian countries (8 countries), Europe (5) and North America (2). The distribution of viral clades as reported from different countries during this time period is summarized in Supplementary Figure 2. Analysis of data on 6181 sequences generated during the past three months revealed that 3789 mutation events distributed over 3772 nucleotide sites. Among these sites, variants at only 11 sites, of which 8 were coding (ORF8 -L84S, ORF1a – V378I, ORF1a – L3606F, ORF1a – A3220V, ORF3a – G251V, ORF1a – L3606F, S – D614G, ORF1b – P314L), were present in relatively high frequencies in multiple populations. These 11 sites were the most useful in defining the phylogenetic clade structure of the viral sequences [Supplementary Figure 3, Supplementary Table 1]. The radial phylogenetic timetree is depicted in Supplementary Figure 1(a), with concentric circles showing the dates of sequence deposition/collection; earlier dates of deposition/collection are closer to the centre. We found that 5629 (91%) of 6181 of SARS-CoV-2 RNA sequences were submitted from East Asian, European and North American regions. The remaining 9% of the sequences were from Oceania, South America, Africa and South-West-Central Asian countries. During 15^th^ January to 31^st^ March 2020 (10 weeks), SARS-CoV-2 had evolved into 11 clades. Four of these 11 clades have attained frequencies higher than 5%; A2a=60.95%, O=13.3%, B1=9% and A1a=7.8% [Supplementary Table 3]. The A2a clade with the highest frequency is defined by nucleotide changes at two “highly informative” sites that are in complete non-random association (linkage disequilibrium): a non-synonymous D614G (Aspartate (D) -> Glycine (G)) mutation in the Spike glycoprotein and a P314L mutation in Orf1b protein of SARS-CoV-2. Of the 11 clades, only 2 clades (a minor A2 subtype and the major A2a subtype) have Glycine at the 614^th^ amino acid position; the remaining 9 clades (ancestral O and evolved clades B, B1, B2, B4, A3, A6, A7 and A1a) have Aspartate [Supplementary Figure 1(b)]. analysis has shown that the most frequent A2a subtype arose in China in mid-January 2020 (Inferred date: January 15^th^, CI: 4^th^-20^th^ January) and spread to multiple locations within 15 days. The initial A2a sequences were deposited from China (Zhejiang/HZ103/2020 on 24^th^ January), and then from another province Shanghai 300Kms away (Shanghai/SH0014/2020 on 28^th^ January and Shanghai/SH0086/2020 on 31^st^ January). The earliest evidence of A2a in Europe was from Germany (Germany/BavPat1/2020 sequence deposited on 28^th^ January). By the end of January, 98.3% of all submitted sequences from East Asia, 90% of all in Europe and 100% of all in North America were of non-A2a subtypes with 614D. The A2a (614G) subtype was then present at a very low frequency in East Asia (3 sequences), Europe (1 sequence) and absent in North America. Dramatic changes in the viral landscape took place very rapidly. By the end of February, the frequency of A2a rose from 9% to 56.25% in Europe and from 0% to 9.7% in North America. Contrastingly, the A2a landscape remained low in China (<1% in February) and other East Asian countries (only 1 A2a sequence out of 206 sequences deposited by the end of February); Figure 1 and Supplementary Figure 4. The A2a – 614G subtype continued to rise in frequency replacing all previously frequent non-A2a-614D subtype in Europe (69% in March comprising 2373 sequences) and in North-America (61% in March comprising 933 sequences) to become the most dominant subtype [Figure 1]. Sequence submission from East Asia dropped drastically in March possibly indicating control of the COVID-19 disease; Supplementary Figure 4. The RNA viruses mutate fast and accumulates changes during transmission process from one infected individual to another. Interestingly, during January 15^th^ and March 31^st^, accumulation of new isolate-specific mutations, were substantially fewer for the A2a clade than for the other clades [Supplementary Figure 5]. These facts are consistent with our negative estimates of Tajima’s D that indicate rapid population expansion coupled with selective sweep of non-A2a-614D clades in East Asia and of A2a-614G clade in Europe and North America [Supplementary Table 4]. Further, the impact of positive selection on A2a may have been higher in Europe than in East Asia, as evidenced by the A2a isolates in Europe and North America having acquired a fewer number of new mutations during this time period than those in East Asia [Supplementary Figure 6].

**Figure 1.**
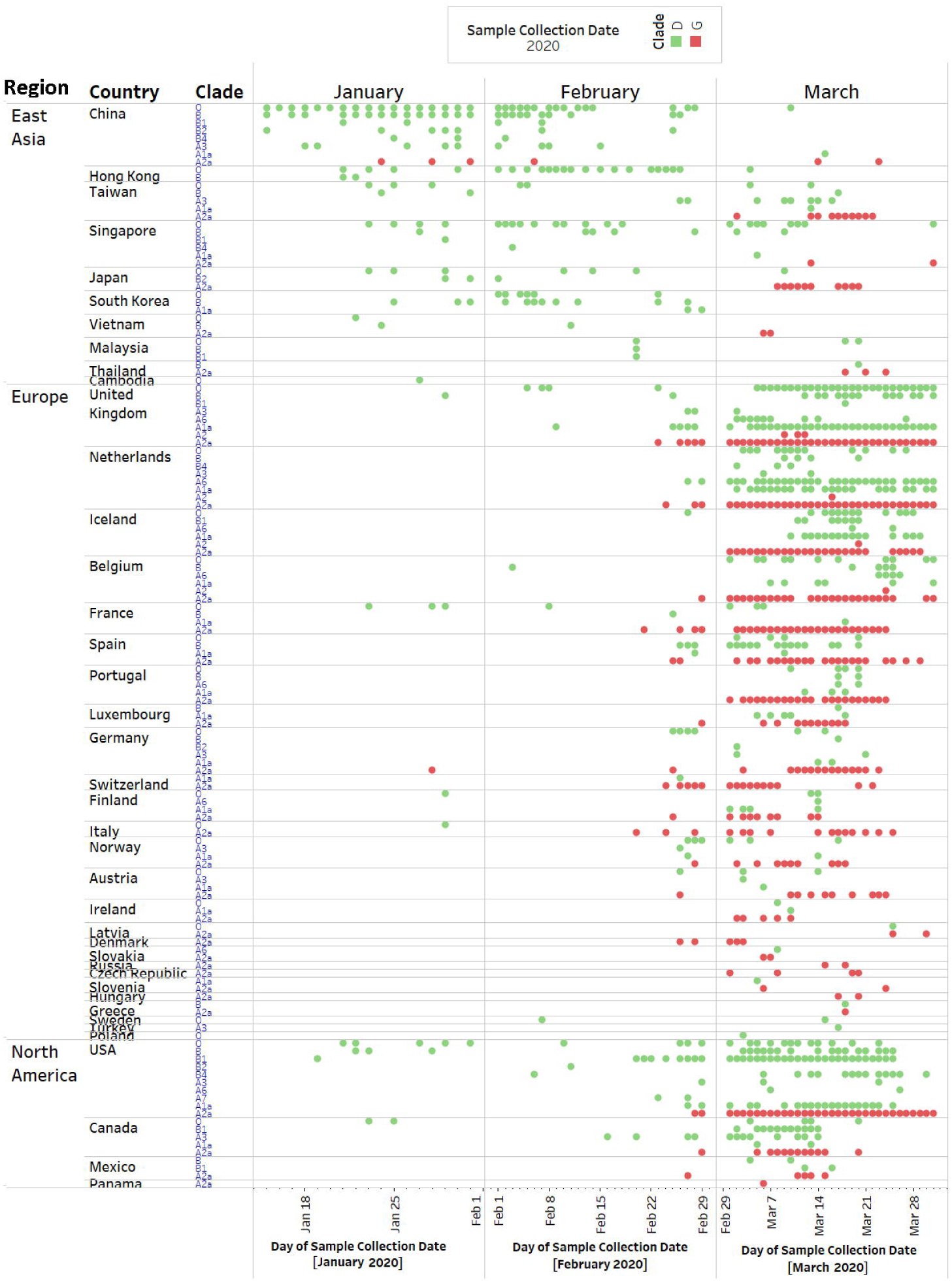
Temporal sample collection pattern for SARS-CoV-2 subtypes from different countries. Day wise sample collection data points for different viral subtypes from individual countries were plotted. The points are color coded based on the presence of D or G (aa) residue in 614 (aa) position of SARS-CoV-2 spike (S) protein.

### The clade specific D614G mutation provides advantage to A2a subtype for host cell entry

The non-synonymous D614G mutation, which defines the A2a clade of SARS-CoV-2, is located between S1-RBD and S2 junctions of the SARS-CoV-2 spike (S) protein [Figure 2 (b)]. The amino acid position 614 is monomorphic for D (Aspartate) residue in SARS-CoVs obtained from bat, civet, pangolin and human [Figure 2 (a)]. It is intriguing that a mutated allele (614G, present in the A2a subtype) at this conserved amino acid site should rapidly rise to a high frequency in some, but not all, regions of the world. *In vitro* introduction of new proteolytic sites at and around the S1-S2 junction is known to substantially increase SARS-CoV fusion with cell membrane (*38*). We have predicted, by proteolytic cleavage prediction analysis, that a novel serine protease (elastase) cleavage site has been introduced in SARS-CoV-2 at 615-616 residues on S protein [Figure 2 (c); Supplementary Table 5]. The position 614 on the S protein is the nearest substrate site for serine protease to cleave at 615-616. The S glycoprotein must be cleaved by host proteases to enable fusion of the viral envelope with the host cell membrane. This fusion is essential for viral entry. The introduction of the new cleavage site for the elastase protease in A2a possibly provides this subtype an advantage over the other subtypes for entry into the host cell. Elastase is mainly expressed by host neutrophils that is elevated in COVID-19 patients (*48–51*). Host TMPRSS2 is a key protein that enables cleavage of the S protein at S1-S2 junction sites (*18, 24*). By gaining an additional elastase cut site, the A2a – 614G subtype is likely to obtain a substantial advantage to enter host cells more efficiently due to simultaneous processing of exogenous elastase proteases and membrane bound TMPRSS2. Of relevance is the fact that SARS-CoV infection was shown to be enhanced by exogenous proteases, including elastase (*52*).

**Figure 2.**
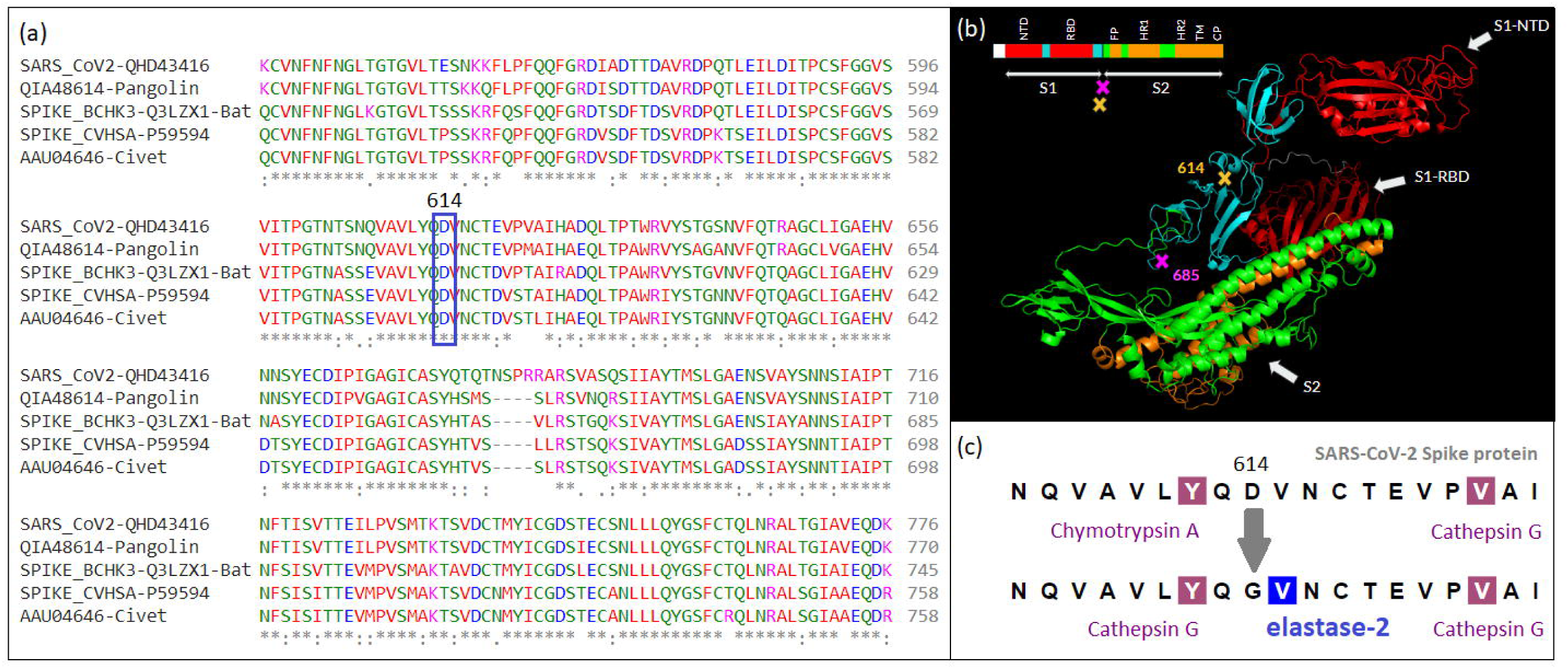
Multiple alignment of spike (S) protein of corona viruses from Bat, Pangolin, Civet and Human SARS-CoV with the Human SARS-CoV-2 showed >70% sequence identity. The 614 amino acid position in the S protein was found to be conserved among species, until the A2a subtype defining mutation occurred. (b) Domain structure of SARS-CoV-2 S protein. Amino acid residues 614 and 685 (S1-S2 junction) are marked on the 3D protein structure. (c) An additional serine protease (elastase) cleavage site around S1-S2 junction was introduced due to D614G mutation in SARS-CoV-2. The Glycine at 614 (aa) is predicted to be the nearest substrate site for elastase to perform proteolytic cleavage in the adjacent residue.

### A common variant at a lung-specific eQTL of *TMPRSS2* and *MX1* accelerated the spread of A2a subtype

The introduction of a novel serine protease (elastase) cleavage site on the S protein resulting from the aspartic acid (D) to glycine (G) mutation at the 614th amino acid position is not completely adequate to explain the rapid rise in A2a frequency [Figure 1]. In order to gain a deeper understanding of this phenomenon, we examined whether there is population genomic variation in *TMPRSS2* and other relevant genomic regions associated with the observed geographical differences in profiles of infection by the subtypes of SARS-CoV-2. We identified 136 eQTLs that regulate expression of *TMPRSS2* in the lung, all of which are located in the 3’ regulatory region of *TMPRSS2*. Of these 136 loci, 64 are also eQTLs for a gene, *MX1* (MX Dynamin Like GTPase 1), which is adjacent to *TMPRSS2* and oriented in the opposite direction. The normalized effect sizes of the non-reference (variant) alleles at these 64 loci on expression levels of *TMPRSS2* and *MX1* are highly correlated (r^2^ = 0.9, p < 2×10^−16^). Variant alleles at 12 of these 64 eQTLs increased the expression of *TMPRSS2* (NES > 0.1) and also of *MX1* (NES > 0.25).

These 12 loci exhibited high levels of differentiation among populations, with F_st_ values exceeding 0.3 for both East Asian – European and East Asian – North American comparisons [Figure 3]. One of these eQTLs (rs35074065), located in the shared 3’ regulatory region of *TMPRSS2* and *MX1*, exhibits the strongest association with increased expression of *TMPRSS2*. The reference allele at rs35074065 is C and the variant is a deletion (delC) of this nucleotide. The delC allele significantly increases the expression of *TMPRSS2* (Normalized Effect Size [NES] of 0.13 (p = 3.9×10^−11^) and *MX1* (NES of 0.2 (p = 1.0×10^−5^) in the lung tissue.

**Figure 3.**
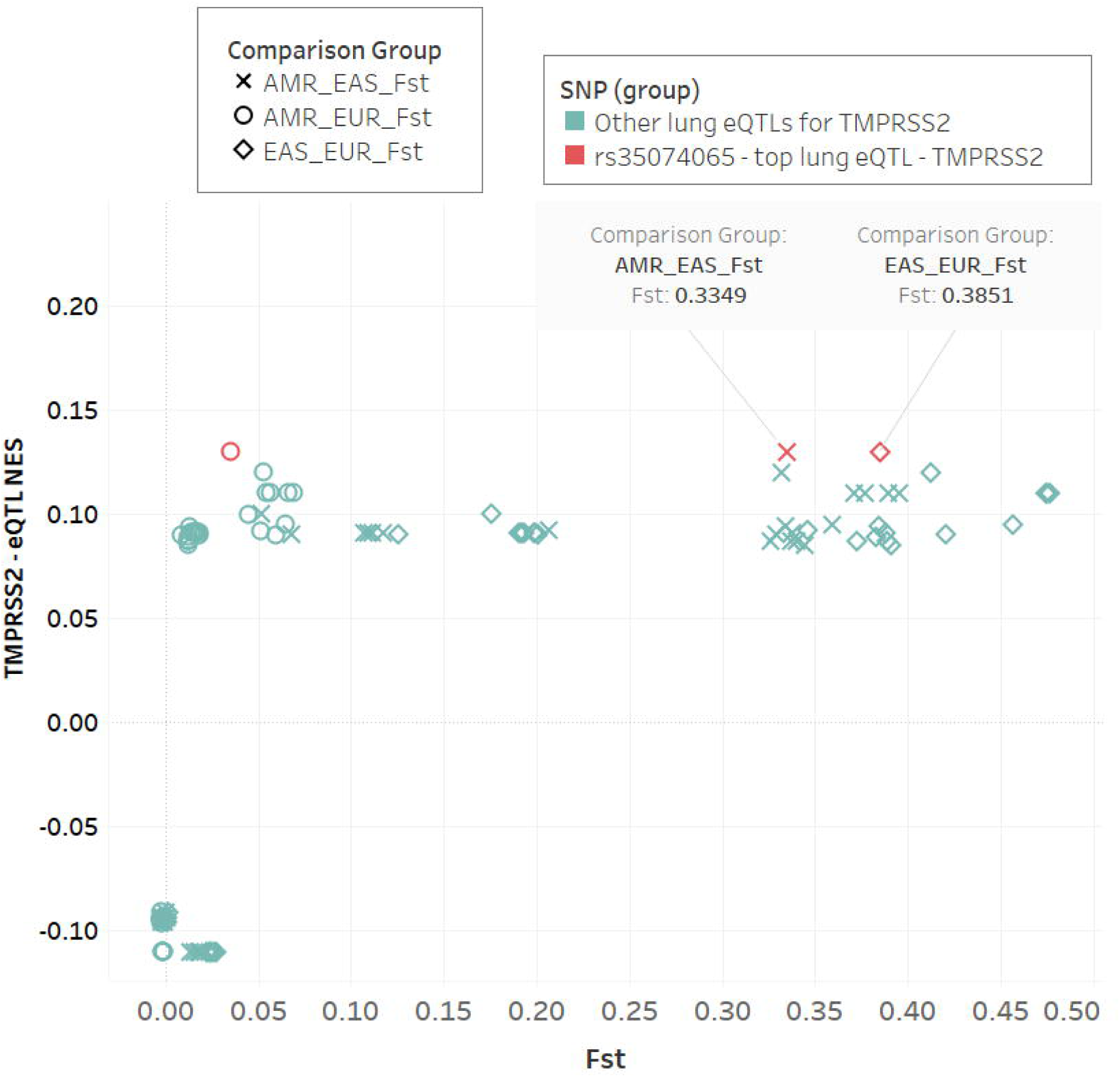
F_st_ across populations for lung-specific TMPRSS2 eQTLs with respective NES. A high F_st_ value between AMR-EAS and EAS-EUR is observed for rs35074065 eQTL with the highest NES for TMPRSS2 gene.

We bioinformatically identified transcription factor binding sites around the rs35074065 locus. A recognition site for transcription factor IRF1 (Interferon Regulatory Factor 1), which is known to activate *MX1* (*53*), was present in the sequences – containing the reference (C) and the variant (delC) alleles – around the rs35074065 eQTL. *IRF1* is known to activate genes in response to viruses (*54*). A transcriptional repressor, IRF2, competes with IRF1 for DNA binding thereby limiting the IRF1 driven pro-inflammatory cascade (*55–57*). The recognition site of transcriptional repressor *IRF2*, which acts antagonistically with *IRF1* (*55, 56, 58*), is absent in the variant (delC) allele containing sequence, but not in the reference (C) allele containing sequence. Loss of the repressor IRF2 recognition site in individuals with the variant (delC) allele is, therefore, expected to further enhance the expression of both *TMPRSS2* and *MX1* genes in the presence of the activator IRF1. The antiviral *MX1* gene activates a cascade of pro-inflammatory effectors (*59, 60*) which leads to neutrophil infiltration (*61–63*). Neutrophil to leukocyte ratio is found to be higher among COVID-19 patients (*48, 49*). Neutrophils are known to secrete the serine protease elastase which has an additional cut site at position 614 for the A2a viral subtype.

The eQTL rs35074065 is highly polymorphic and exhibits wide variation in allele frequencies across continental populations. The delC allele is rare among East-Asians (allele frequency: 0.0124), occurs at high frequencies among Europeans (∼0.4), and at intermediate frequencies among North Americans (0.26) [Figure 4]. We have found the delC allele frequency to strongly correlate (r^2^ = 0.91) with the frequency of the A2a subtype across geographical regions [Supplementary figure 7].

**Figure 4.**
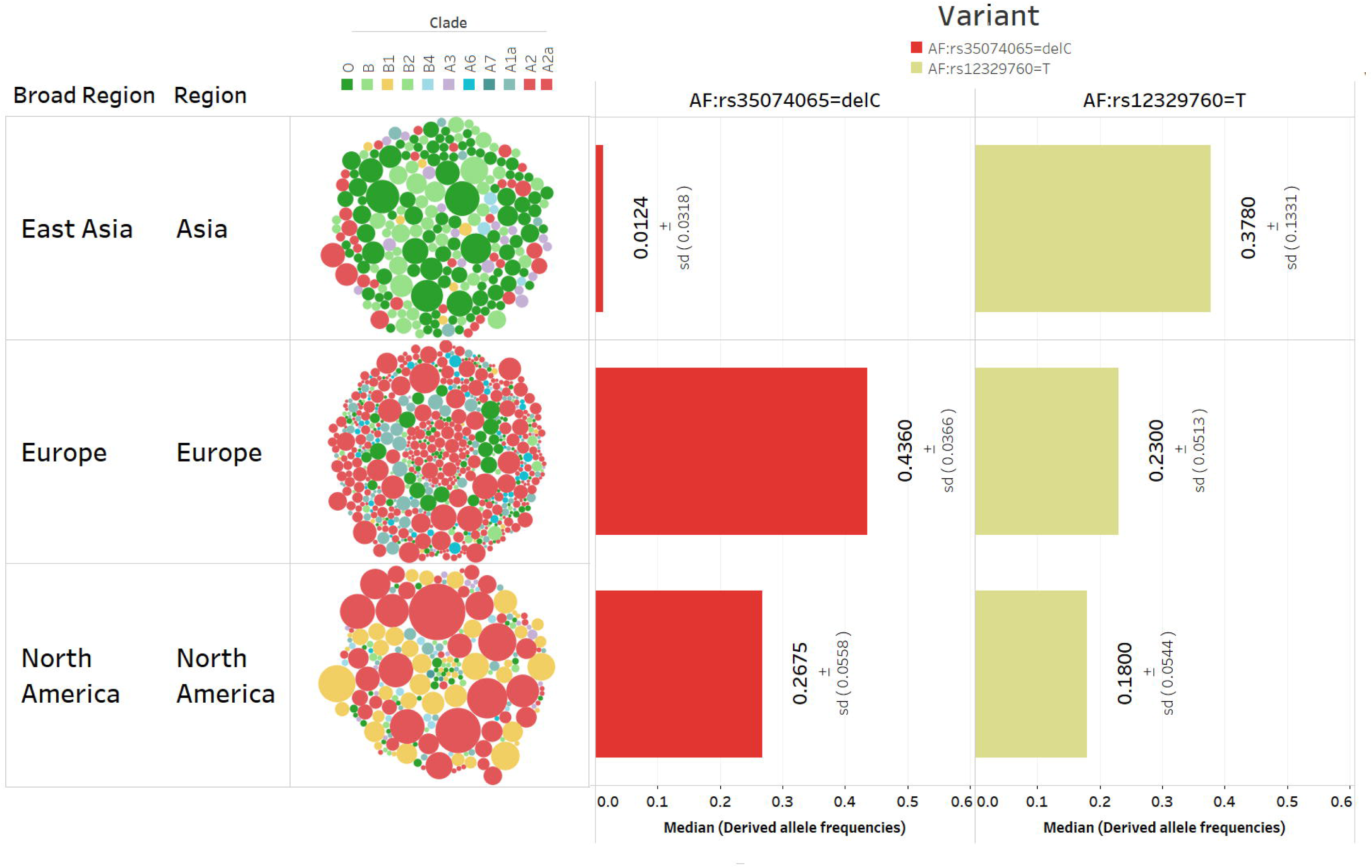
(a) SARS-CoV-2 subtype distribution in different populations. Circle size is proportional to the number of sequences submitted on a particular date and the circle colour distinguishes the subtypes. The positions of circles from the centre towards the outer periphery depicts the date of submission from January to March [more recent submissions are towards the periphery]. (b) The vertical bars represent median derived allele frequencies of *TMPRSS2* and *MX1* eQTL (rs35074065:del-C) and TMPRSS2 missense polymorphic variant (rs12329760:T), respectively, for different continental populations (East Asia = 13 populations, Europe = 11, North America = 5). The standard deviation of the derived allele frequencies are provided within parentheses.

### A highly polymorphic *TMPRSS2* coding variant protects East Asians from infection by A2a subtype

As we have noted, the proportion of infected persons with the A2a subtype is rather low in East Asia, but high in Europe and North America. We postulated that non-silent variants in *TMPRSS2* that are common among East Asians, but not among Europeans or North Americans, may confer protection to them against being infected by A2a. We have identified 16 non-silent coding variants (details with predictive scores of function in Supplementary Table 6) from the 1000Genomes database. Two non-silent variants (rs12329760 and rs75603675), of which one (rs12329760) is in an evolutionarily conserved region, showed high allele frequency differences between East Asians and other populations [Supplementary Table 7].

The coding missense variant rs12329760 – V160M, but not rs75603675, is likely pathogenic, since it is predicted to be highly “Deleterious” by SIFT (score = 0.01), Polyphen2, (score = 0.9), FATHMM-MKL (score = 0.76) and CADD (score = 28.5). Further, it was identified that rs12329760, with a deleterious variant allele T, is not in linkage disequilibrium in any population (r^2^; European = 0.1, North American = 0.03, East Asian = 0.001) with the rs35074065 – delC eQTL locus, which is at an 18.9 Kb physical distance.

The deleterious SNP (rs12329760) resides within the extracellular highly conserved SRCR (scavenger receptor cysteine-rich) domain of *TMPRSS2* (*64*). SRCR domains are known to interact with external pathogens (*65*). Functional studies have shown that the proteolytic activity driven by the serine protease domain of TMPRSS2 reduces drastically without the help of SRCR domain (*66*). The SRCR domain functions to properly orient the protease domain towards its substrate (*64*). Based on these facts, it appears that this nonsynonymous deleterious coding variant of *TMPRSS2* on SRCR domain may have altered activity for processing the Spike protein of SARS-CoV-2 A2a subtype which has the 614G mutation, in an evolutionary conserved region. We have observed that the frequency of rs12329760-T variant allele is negatively correlated (r^2^ = −0.4) with proportion of A2a type frequency [Supplementary Figure 8]. Among East Asians, individuals with TT genotype (∼19%), may have a genetic protective advantage against A2a viral type; compared to North Americans (TT = ∼4%) and Europeans (TT = ∼7%) populations.

## Discussion

Within four months, the COVID-19 pandemic spread rapidly to more than 200 countries in different continents. SARS-CoV-2 has shown a strong association with increased morbidity and mortality in specific countries, notably USA and Western European countries. We sought to understand whether the observed geographical variations in prevalence of infection by distinct SARS-CoV-2 subtypes can be explained, at least in part, by variation in specific genes of the human host populations.

We performed analysis on publicly available data of RNA sequences of SARS-Cov-2 isolates to correlate pathogen evolution with disease transmission. We have shown that a particular subtype −A2a − that rose in frequency in East Asia in January 2020, has spread rapidly through the European and North American continents. The spread of A2a has been so explosive, that in 10 weeks (of February and March 2020) over 60% of humans were infected with A2a starting from only 2%. The A2a subtype acquired few mutations during the transmission, even lesser in Europe [Supplementary Figure 6].

The A2a subtype is characterized by the D614G mutation near the S1-S2 junction on the Spike protein. D614G mutation introduces an additional cleavage site in the S protein that is specific for elastase, a serine protease, generally produced by the neutrophils. Sequential digestion at multiple sites on the S1-S2 junction of S protein is necessary for SARS-CoV-2, to enter into the host cell (*21, 38*). The cascade of cleavage is also dependent on the availability of host protease in the vicinity (*38, 52*). Multiple digestions at the S1-S2 junction region produce “decoy” fragments which reduces host immune response by binding and inactivating antiviral antibodies (*67*) giving the virion an advantage to overcome the first line of host defence. It has been shown that for cellular entry of SARS-CoV-2, ACE2 and TMPRSS2 are indispensable, while endosomal cysteine proteases cathepsin B and L (CatB/L) are not (*24, 28*).

Our analysis showed that protein altering missense variants are rare (MAF < 0.01) in the genes of relevance to the infection, including ACE2, Cathepsin-L, MX1 and elastase [Supplementary Table 8-11]. However, another gene of great relevance, *TMPRSS2*, harbors variants that are common in all global populations. Therefore, we focused on genetic variations in *TMPRSS2* to explore the role of host genomic variants in modulating susceptibility to infection.

We identified lung-specific eQTL (rs35074065 [C > – (del)]) that has a major regulatory control of the expression of *TMPRSS2*. This eQTL showed significant allele frequency differences (high F_st_) between East Asians (Median derived allele frequencies [DAF] all sub populations: 0.01; sd: 0.030) and Europeans (Median DAF: 0.44; sd: 0.04) and Americans (Median DAF: 0.27; sd: 0.06), as estimated from 29 global populations residing in these regions [Figure 4]. This eQTL also regulates the expression of *TMPRSS2* as well as *MX1*, a nearby gene with an opposite orientation to *TMPRSS2. MX1* modulates the type-I interferon mediated anti-viral inflammatory response in lungs (*68*). Our transcription factor binding site prediction analysis identified that both IRF1 transcription factor and IRF2 repressor bind to the reference allele (C) of rs35074065. The rs35074065-delC common variant abolishes the recognition site for the repressor IRF2, but not of the transcription factor IRF1. IRF1 and IRF2 are antagonistic to each other (*55, 56, 58*), but help maintain a fine balance in expression of both *MX1* and *TMPRSS2*. Loss of IRF2 recognition site, due to rs35074065-delC variant, leads to increased IRF1 mediated expression of *MX1* (*53*). Recent analysis by Human Lung Cell Atlas group identified that *ACE2* and *TMPRSS2* are interferon regulated and ACE2+ TMPRSS2+ Single cells that express both ACE2 and TMPRSS2, also express innate immunity associated genes like *MX1, IDO1*, etc. (*18*) Increased *MX1* expression is known to direct inflammatory responses via IFN type-I activation (*59, 61, 68*) which in turn mediates neutrophil infiltration (*61, 69–72*). Neutrophils are known to express elastase proteins at a very high level. The spike protein of SARS-CoV-2 subtype A2a has an additional elastase specific proteolytic cleavage site for 614G amino acid. A2a, therefore, enjoys a double advantage in its ability to gain entry into human hosts in Europe and North America: (1) A2a – 614G has an additional cleavage site in the S protein that is specific for elastase protease, which is absent in the 614D subtypes (2) European and North American populations have high frequencies of the delC allele at rs35074065 eQTL, which is associated with increased expression of TMPRSS2 [Figure 5]. A recent single cell ATAC seq study also reported that eQTL-rs35074065 acts as transcriptional enhancer and showed increased expression of *TMPRSS2* in lung tissues (*73*). The simultaneous processing of S protein by elastase and TMPRSS2 proteases help the A2a subtype coronaviruses to more easily infect the large majority of Europeans and North Americans who possess the delC allele [Figure 5]. Elevated levels of neutrophils are reported in SARS-CoV-2 infected patients with increased disease severity (*48–51*). Our findings are indicative of the possibility of using elastase inhibitors as therapy to prevent infection by the A2a subtype of SARS-CoV-2. We note that the use of elastase inhibitors in chronic obstructive pulmonary disease is under active consideration (*74, 75*).

**Figure 5.**
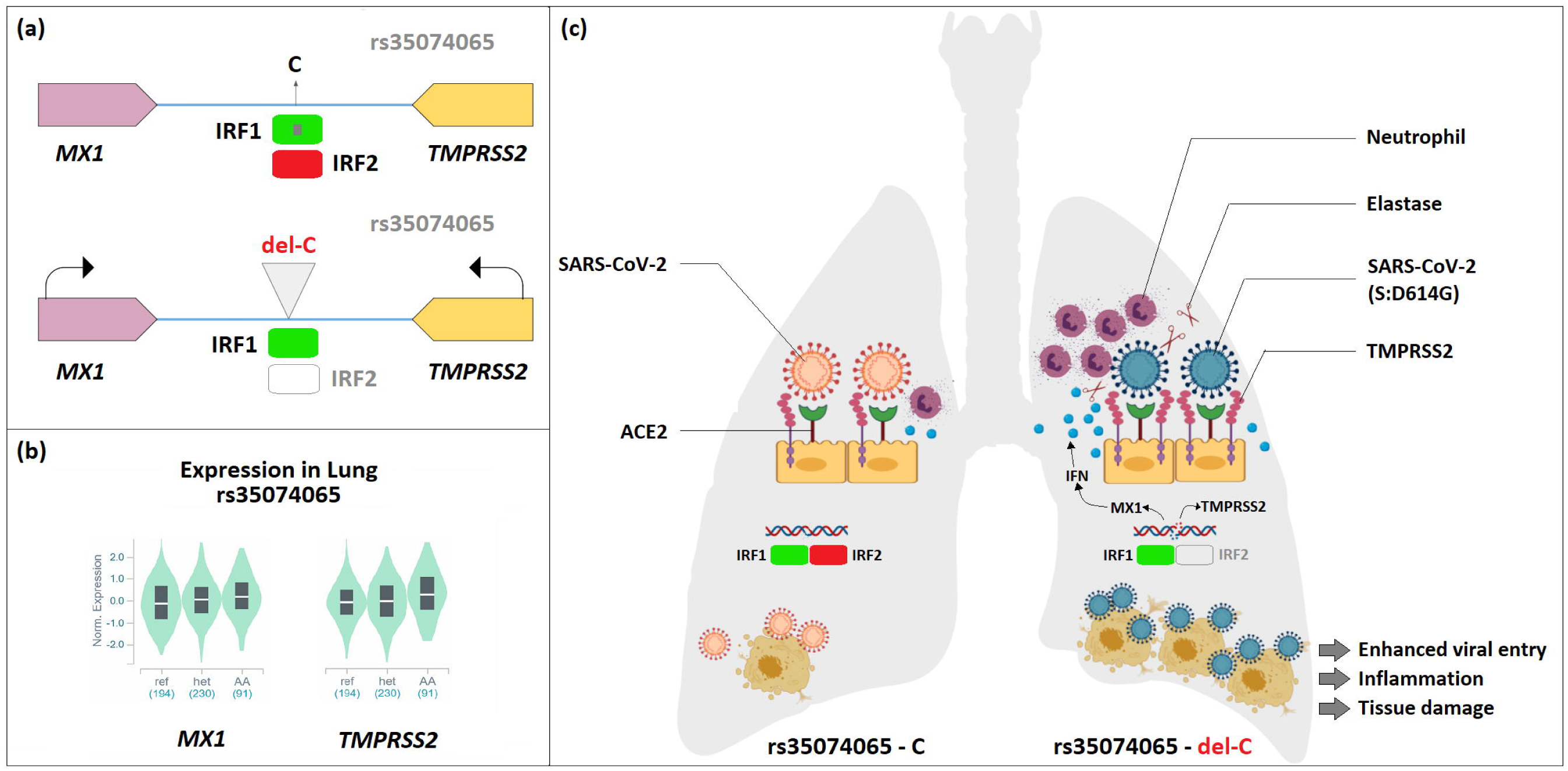
(a) Gene expression regulation of TMPRSS2 and MX1 by an interplay of transcriptional enhancer IRF1 and repressor IRF2. Loss of IRF2 recognition site due to deletion of C at rs35074065 enhances IRF1 mediated TMPRSS2 and MX1 expression. (b) Expression of TMPRSS2 and MX1 in the lung for reference allele homozygous, heterozygous, and variant allele homozygous genotypes at rs35074065. (c) The upregulation of TMPRSS2 enhances proteolytic cleavage of SARS-CoV-2 S protein and entry into lung cell of host. MX1 upregulation results in IFN type-I mediated neutrophil infiltration in the lung. With elevated levels of TMPRSS2 and neutrophil Elastase, SARS-CoV-2 A2a-614G subtype having additional Elastase specific cut site near S1-S2 junction facilitates dual processing by host serine proteases, TMPRSS2 and Elastase (ELANE).

We have also identified a non-silent damaging variant in a highly conserved SRCR domain (rs12329760; V160M) of *TMPRSS2* that is present in East Asians at double the frequency (MAF=0.4) than in Europeans (MAF=0.2). The SRCR domain interacts with external pathogens (*65, 76, 77*) and helps in positioning of the protease head towards the substrate (*64*). The SARS-CoV-2 A2a subtype specific D614G mutation is in a region of the S protein that is highly conserved across the coronavirus family. We predict that the V160M mutation in the host and D614G mutation in the A2a subtype will result in reduced interaction of the SRCR domain of TMPRSS2 and the S protein of the coronavirus leading to a reduction in viral entry in the host cell. High prevalence of the V160M mutation in populations of East Asia is a likely reason for the reduced spread of A2a subtype in East Asia.

In this study, we have provided strong evidence that host genetic variation regulating the expression of the *TMPRSS2* gene has shaped the global spread of the A2a subtype of SARS-CoV-2.

## Acknowledgment

All viral sequence data related to SARS-Cov-2 was available from Global Initiative on Sharing All Influenza Data (GISAID) EpiFlu database and NIH Genbank resource. We acknowledge all the contributions of novel coronavirus sequencing data by members of the broader scientific community and GISAID for making everything accessible. The credits for sample originating and sequence submitting lab details are provided in Supplementary Table 12. We are also grateful to Prof. Sharmila Sengupta and Dr Indranil Banerjee for sharing some valuable comments on the manuscript.

## Conflicts of Interest

None

## Author Contributions

N.K.B conceived the study. P.P.M, A.B, S.M and N.K.B shaped up the study design. C.B, C.D, A.G., A.K.S, and N.K.B involved in data mining and participated in data analysis. A.K.S, C.D and N.K.B performed virus epidemiological analyses. C.B, C.D, A.G and N.K.B analysed human genomic data. A.G and C.D performed structural modelling and TFBS analysis. All authors were involved in manuscript writing. A.B and P.P.M edited the final manuscript. All authors read and approved the contents of the manuscript.

## LEGENDS TO FIGURES

**Supplementary Figure 1**.

Frequency distribution of viral subtypes (clades) in different countries during January 15 and March 31, 2020

**Supplementary Figure 2**.

(a) Nucleotide diversity and (b) amino acid diversity in 6181 SARS-CoV-2 genomes collected worldwide from 62 countries. Only 8 coding sites, i.e. ORF8 -L84S, ORF1a – V378I, ORF1a – L3606F, ORF1a – A3220V, ORF3a – G251V, ORF1a – L3606F, S – D614G, ORF1b – P314L and 3 other nucleotide sites were present in reasonably high frequencies. The viral clade/subtype in which these mutations are present are shown at the bottom panel of the figure.

**Supplementary Figure 3**.

(a) Radial phylogenetic timetree based on 6181 SARS-CoV-2 RNA sequences that comprise 11 clades. The concentric circles depict the date of sequence deposition/collection; earlier date of deposition/collection is closer to the centre. (b) The radial phylogenetic time tree is annotated by presence of D or G residue at 614th amino acid position.

**Supplementary Figure 4**.

Frequency distributions of SARS-CoV-2 subtypes 614D and 614G in East Asia, Europe and North America for the months of January, February and March 2020. The size of the circles represent the number of samples belonging to a specific subtype collected on a specific date on the month. The circles are color coded based on the presence of D or G amino acid in 614 position of S protein. The circles close to the center belong to early sequence deposition of a specific month and circles in the boundaries are late submission.

**Supplementary Figure 5**.

The number of isolate specific mutations acquired by different SARS-CoV-2 subtypes during 15th January to 31st March, 2020. The clades are color coded as follows: 614D in blue and 614G-A2a clade in red). The A2a specific sequences have acquired less isolate specific mutations over time.

**Supplementary Figure 6**.

Number of isolate specific mutations in populations of different regions acquired by SARS-CoV-2 subtypes during 15th January to 31st March, 2020), separately for D and G amino acids at 614 amino acid position on spike (S) protein. The clades are color coded as follows: 614D in blue and 614G-A2a clade in red. The 614G (A2a) isolates in Europe have acquired a smaller number of mutations compared to East Asian A2a-614G isolates during the same time period.

**Supplementary Figure 7**.

Correlation (r2 = 0.9) of eQTL-rs35074065:del-C allele frequency with the SARS-CoV-2 A2a subtype frequencies (averaged of frequencies from 15th January to 31st March) in East Asia, Europe and America.

**Supplementary Figure 8**.

Correlation (r2 = −0.4) of TMPRSS2 coding SNP rs12329760:T allele frequency with the SARS-CoV-2 A2a subtype frequencies (averaged of frequencies from 15th January to 31st March) in East Asia, Europe and America.

## Notes

### Competing Interest Statement

The authors have declared no competing interest.

## References

1. R. Verity, L. C. Okell, I. Dorigatti, P. Winskill, C. Whittaker, N. Imai, G. Cuomo-Dannenburg, H. Thompson, P. G. T. Walker, H. Fu, A. Dighe, J. T. Griffin, M. Baguelin, S. Bhatia, A. Boonyasiri, A. Cori, Z. Cucunubá, R. FitzJohn, K. Gaythorpe, W. Green, A. Hamlet, W. Hinsley, D. Laydon, G. Nedjati-Gilani, S. Riley, S. van Elsland, E. Volz, H. Wang, Y. Wang, X. Xi, C. A. Donnelly, A. C. Ghani, N. M. Ferguson, Estimates of the severity of coronavirus disease 2019: a model-based analysis. Lancet. Infect. Dis. (2020), doi: 10.1016/S1473-3099(20)30243-7.

2. N. Zhu, D. Zhang, W. Wang, X. Li, B. Yang, J. Song, X. Zhao, B. Huang, W. Shi, R. Lu, P. Niu, F. Zhan, X. Ma, D. Wang, W. Xu, G. Wu, G. F. Gao, W. Tan, A novel coronavirus from patients with pneumonia in China, 2019. N. Engl. J. Med. 382, 727–733 (2020).

3. A. E. Gorbalenya, S. C. Baker, R. S. Baric, R. J. de Groot, C. Drosten, A. A. Gulyaeva, B. L. Haagmans, C. Lauber, A. M. Leontovich, B. W. Neuman, D. Penzar, S. Perlman, L. L. M. Poon, D. V. Samborskiy, I. A. Sidorov, I. Sola, J. Ziebuhr, The species Severe acute respiratory syndrome-related coronavirus: classifying 2019-nCoV and naming it SARS-CoV-2. Nat. Microbiol. 5, 536–544 (2020).

4. Y. Chen, L. Li, SARS-CoV-2: virus dynamics and host response. Lancet Infect. Dis. 20, 515–516 (2020).

5. WHO, WHO Director-General’s opening remarks at the media briefing on COVID-19 – 11 March 2020 (2020), (available at <https://www.who.int/dg/speeches/detail/who-director-general-s-opening-remarks-at-the-media-briefing-on-covid-19---11-march-2020).

6. WHO, Coronavirus Disease 2019 (COVID-19) Situation Reports. April 1 2020. WHO Situat. Rep. 2019 (2020), pp. 1–19.

7. C. Wang, P. W. Horby, F. G. Hayden, G. F. Gao, A novel coronavirus outbreak of global health concern. Lancet. 395, 470–473 (2020).

8. J. Sun, W. T. He, L. Wang, A. Lai, X. Ji, X. Zhai, G. Li, M. A. Suchard, J. Tian, J. Zhou, M. Veit, S. Su, COVID-19: Epidemiology, Evolution, and Cross-Disciplinary Perspectives. Trends Mol. Med. (2020), doi: 10.1016/j.molmed.2020.02.008.

9. CDC COVID-19 Response Team, Coronavirus Disease 2019 in Children – United States, February 12-April 2, 2020. MMWR. Morb. Mortal. Wkly. Rep. 69, 422–426 (2020).

10. P. Spychalski, A. Blazynska-Spychalska, J. Kobiela, Estimating case fatality rates of COVID-19. Lancet Infect. Dis. (2020), doi: 10.1016/S1473-3099(20)30246-2.

11. M. A. Khafaie, F. Rahim, Cross-country comparison of case fatality rates of Covid-19/SARS-CoV-2. Osong Public Heal. Res. Perspect. 11, 74–80 (2020).

12. G. Onder, G. Rezza, S. Brusaferro, Case-Fatality Rate and Characteristics of Patients Dying in Relation to COVID-19 in Italy. JAMA (2020), doi: 10.1001/jama.2020.4683.

13. N. K. Biswas, P. P. Majumder, Analysis of RNA Sequences of 3636 SARS-CoV-2 Collected from 55 Countries Reveals Selective Sweep of One Virus Type. Indian J. Med. Res. (ACCEPTED) (2020).

14. P. Forster, L. Forster, C. Renfrew, M. Forster, Phylogenetic network analysis of SARS-CoV-2 genomes. Proc. Natl. Acad. Sci. 117, 202004999 (2020).

15. J. Hadfield, C. Megill, S. M. Bell, J. Huddleston, B. Potter, C. Callender, P. Sagulenko, T. Bedford, R. A. Neher, NextStrain: Real-time tracking of pathogen evolution. Bioinformatics. 34, 4121–4123 (2018).

16. D. F. Gudbjartsson, A. Helgason, H. Jonsson, O. T. Magnusson, P. Melsted, G. L. Norddahl, J. Saemundsdottir, A. Sigurdsson, P. Sulem, A. B. Agustsdottir, B. Eiriksdottir, R. Fridriksdottir, E. E. Gardarsdottir, G. Georgsson, O. S. Gretarsdottir, K. R. Gudmundsson, T. R. Gunnarsdottir, A. Gylfason, H. Holm, B. O. Jensson, A. Jonasdottir, F. Jonsson, K. S. Josefsdottir, T. Kristjansson, D. N. Magnusdottir, L. le Roux, G. Sigmundsdottir, G. Sveinbjornsson, K. E. Sveinsdottir, M. Sveinsdottir, E. A. Thorarensen, B. Thorbjornsson, A. Löve, G. Masson, I. Jonsdottir, A. D. Möller, T. Gudnason, K. G. Kristinsson, U. Thorsteinsdottir, K. Stefansson, Spread of SARS-CoV-2 in the Icelandic Population. N. Engl. J. Med. (2020), doi: 10.1056/nejmoa2006100.

17. P. Yang, X. Wang, COVID-19: a new challenge for human beings. Cell. Mol. Immunol. (2020), doi: 10.1038/s41423-020-0407-x.

18. C. Muus, M. D. Luecken, G. Eraslan, A. Waghray, G. Heimberg, L. Sikkema, Y. Kobayashi, E. D. Vaishnav, A. Subramanian, C. Smilie, K. Jagadeesh, E. T. Duong, E. Fiskin, E. T. Triglia, M. Ansari, P. Cai, B. Lin, J. Buchanan, S. Chen, J. Shu, A. L. Haber, H. Chung, D. T. Montoro, T. Adams, H. Aliee, J. Samuel, A. Z. Andrusivova, I. Angelidis, O. Ashenberg, K. Bassler, C. Bécavin, I. Benhar, J. Bergenstr\r ahle, L. Bergenstr\r ahle, L. Bolt, E. Braun, L. T. Bui, M. Chaffin, E. Chichelnitskiy, J. Chiou, T. M. Conlon, M. S. Cuoco, M. Deprez, D. S. Fischer, A. Gillich, J. Gould, M. Guo, A. J. Gutierrez, A. C. Habermann, T. Harvey, P. He, X. Hou, L. Hu, A. Jaiswal, P. Jiang, T. Kapellos, C. S. Kuo, L. Larsson, M. A. Leney-Greene, K. Lim, M. Litvi\v nuková, J. Lu, L. S. Ludwig, W. Luo, H. Maatz, E. Madissoon, L. Mamanova, K. Manakongtreecheep, C.-H. Marquette, I. Mbano, A. M. McAdams, R. J. Metzger, A. N. Nabhan, S. K. Nyquist, L. Penland, O. B. Poirion, S. Poli, C. Qi, R. Queen, D. Reichart, I. Rosas, J. Schupp, R. Sinha, R. V Sit, K. Slowikowski, M. Slyper, N. Smith, A. Sountoulidis, M. Strunz, D. Sun, C. Talavera-López, P. Tan, J. Tantivit, K. J. Travaglini, N. R. Tucker, K. Vernon, M. H. Wadsworth, J. Waldman, X. Wang, W. Yan, W. Zhao, C. G. K. Ziegler, Integrated analyses of single-cell atlases reveal age, gender, and smoking status associations with cell type-specific expression of mediators of SARS-CoV-2 viral entry and highlights inflammatory programs in putative target cells. bioRxiv (2020), doi: 10.1101/2020.04.19.049254.

19. W. Wang, Y. Xu, R. Gao, R. Lu, K. Han, G. Wu, W. Tan, Detection of SARS-CoV-2 in Different Types of Clinical Specimens. JAMA (2020), doi: 10.1001/jama.2020.3786.

20. M. M. Lamers, J. Beumer, J. van der Vaart, K. Knoops, J. Puschhof, T. I. Breugem, R. B. G. Ravelli, J. van Schayck, A. Z. Mykytyn, H. Q. Duimel, E. van Donselaar, S. Riesebosch, H. J. H. Kuijpers, D. Schippers, W. J. van de Wetering, M. de Graaf, M. Koopmans, E. Cuppen, P. J. Peters, B. L. Haagmans, H. Clevers, SARS-CoV-2 productively infects human gut enterocytes. Science (80-.). (2020), doi: 10.1126/science.abc1669.

21. A. C. Walls, Y. J. Park, M. A. Tortorici, A. Wall, A. T. McGuire, D. Veesler, Structure, Function, and Antigenicity of the SARS-CoV-2 Spike Glycoprotein. Cell. 181, 281–292.e6 (2020).

22. K. G. Andersen, A. Rambaut, W. I. Lipkin, E. C. Holmes, R. F. Garry, The proximal origin of SARS-CoV-2. Nat. Med. 26, 450–452 (2020).

23. K. Shirato, M. Kawase, S. Matsuyama, Wild-type human coronaviruses prefer cell-surface TMPRSS2 to endosomal cathepsins for cell entry. Virology. 517, 9–15 (2018).

24. M. Hoffmann, H. Kleine-Weber, S. Schroeder, N. Krüger, T. Herrler, S. Erichsen, T. S. Schiergens, G. Herrler, N. H. Wu, A. Nitsche, M. A. Müller, C. Drosten, S. Pöhlmann, SARS-CoV-2 Cell Entry Depends on ACE2 and TMPRSS2 and Is Blocked by a Clinically Proven Protease Inhibitor. Cell. 181, 271–280.e8 (2020).

25. X. X. Qu, P. Hao, X. J. Song, S. M. Jiang, Y. X. Liu, P. G. Wang, X. Rao, H. D. Song, S. Y. Wang, Y. Zuo, A. H. Zheng, M. Luo, H. L. Wang, F. Deng, H. Z. Wang, Z. H. Hu, M. X. Ding, G. P. Zhao, H. K. Deng, Identification of two critical amino acid residues of the severe acute respiratory syndrome coronavirus spike protein for its variation in zoonotic tropism transition via a double substitution strategy. J. Biol. Chem. 280, 29588–29595 (2005).

26. S. Matsuyama, N. Nao, K. Shirato, M. Kawase, S. Saito, I. Takayama, N. Nagata, T. Sekizuka, H. Katoh, F. Kato, M. Sakata, M. Tahara, S. Kutsuna, N. Ohmagari, M. Kuroda, T. Suzuki, T. Kageyama, M. Takeda, Enhanced isolation of SARS-CoV-2 by TMPRSS2-expressing cells. Proc. Natl. Acad. Sci. 117, 7001–7003 (2020).

27. Q. Wang, Y. Zhang, L. Wu, S. Niu, C. Song, Z. Zhang, G. Lu, C. Qiao, Y. Hu, K. Y. Yuen, Q. Wang, H. Zhou, J. Yan, J. Qi, Structural and Functional Basis of SARS-CoV-2 Entry by Using Human ACE2. Cell (2020), doi: 10.1016/j.cell.2020.03.045.

28. S. Xia, L. Yan, W. Xu, A. S. Agrawal, A. Algaissi, C.-T. K. Tseng, Q. Wang, L. Du, W. Tan, I. A. Wilson, S. Jiang, B. Yang, L. Lu, A pan-coronavirus fusion inhibitor targeting the HR1 domain of human coronavirus spike. Sci. Adv. 5 (2019), doi: 10.1126/sciadv.aav4580.

29. E. W. Stawiski, D. Diwanji, K. Suryamohan, R. Gupta, F. A. Fellouse, J. F. Sathirapongsasuti, J. Liu, Y.-P. Jiang, A. Ratan, M. Mis, D. Santhosh, S. Somasekar, S. Mohan, S. Phalke, B. Kuriakose, A. Antony, J. R. Junutula, S. C. Schuster, N. Jura, S. Seshagiri, Human ACE2 receptor polymorphisms predict SARS-CoV-2 susceptibility, bioRxiv, in press, doi: 10.1101/2020.04.07.024752.

30. A. Renieri, E. Benetti, R. Tita, O. Spiga, A. Ciolfi, G. Birolo, A. Bruselles, G. Doddato, A. Giliberti, C. Marconi, F. Musacchia, T. Pippucci, A. Torella, A. Trezza, F. Valentino, M. Baldassarri, A. Brusco, R. Asselta, B. Mirella, S. Furini, M. Seri, V. Nigro, G. Matullo, M. Tartaglia, F. Mari, A. Pinto, ACE2 variants underlie interindividual variability and susceptibility to COVID-19 in Italian population, medRxiv, in press, doi: 10.1101/2020.04.03.20047977.

31. Y. Shu, J. McCauley, GISAID: Global initiative on sharing all influenza data – from vision to reality. Eurosurveillance. 22 (2017),, doi: 10.2807/1560-7917.ES.2017.22.13.30494.

32. E. M. Volz, K. Koelle, T. Bedford, Viral Phylodynamics. PLoS Comput. Biol. 9, e1002947 (2013).

33. K. Katoh, D. M. Standley, MAFFT multiple sequence alignment software version 7: Improvements in performance and usability. Mol. Biol. Evol. 30, 772–780 (2013).

34. L.-T. Nguyen, H. A. Schmidt, A. von Haeseler, B. Q. Minh, IQ-TREE: A Fast and Effective Stochastic Algorithm for Estimating Maximum-Likelihood Phylogenies. Mol. Biol. Evol. 32, 268–274 (2014).

35. F. Tajima, Statistical method for testing the neutral mutation hypothesis by DNA polymorphism. Genetics. 123, 585–595 (1989).

36. S. Kumar, K. Tamura, M. Nei, MEGA: Molecular evolutionary genetics analysis software for microcomputers. Bioinformatics. 10, 189–191 (1994).

37. P. Librado, J. Rozas, DnaSP v5: A software for comprehensive analysis of DNA polymorphism data. Bioinformatics. 25, 1451–1452 (2009).

38. S. Belouzard, V. C. Chu, G. R. Whittaker, Activation of the SARS coronavirus spike protein via sequential proteolytic cleavage at two distinct sites. Proc. Natl. Acad. Sci. U. S. A. 106, 5871–5876 (2009).

39. A. Roy, A. Kucukural, Y. Zhang, I-TASSER: A unified platform for automated protein structure and function prediction. Nat. Protoc. 5, 725–738 (2010).

40. J. Song, H. Tan, A. J. Perry, T. Akutsu, G. I. Webb, J. C. Whisstock, R. N. Pike, PROSPER: An Integrated Feature-Based Tool for Predicting Protease Substrate Cleavage Sites. PLoS One. 7, e50300 (2012).

41. J. Song, F. Li, A. Leier, T. T. Marquez-Lago, T. Akutsu, G. Haffari, K. C. Chou, G. I. Webb, R. N. Pike, PROSPERous: High-throughput prediction of substrate cleavage sites for 90 proteases with improved accuracy. Bioinformatics. 34, 684–687 (2018).

42. L. J. Carithers, H. M. Moore, The Genotype-Tissue Expression (GTEx) Project. Biopreserv. Biobank. 13, 307–308 (2015).

43. H. Li, Tabix: fast retrieval of sequence features from generic TAB-delimited files. Bioinformatics. 27, 718–719 (2011).

44. K. Wang, M. Li, H. Hakonarson, ANNOVAR: Functional annotation of genetic variants from high-throughput sequencing data. Nucleic Acids Res. 38, e164–e164 (2010).

45. B. S. Weir, C. C. Cockerham, Estimating F-Statistics for the Analysis of Population Structure. Evolution (N. Y). 38, 1358–1370 (1984).

46. C. C. Chang, C. C. Chow, L. C. A. M. Tellier, S. Vattikuti, S. M. Purcell, J. J. Lee, Second-generation PLINK: Rising to the challenge of larger and richer datasets. Gigascience. 4 (2015), doi: 10.1186/s13742-015-0047-8.

47. O. Fornes, J. A. Castro-Mondragon, A. Khan, R. van der Lee, X. Zhang, P. A. Richmond, B. P. Modi, S. Correard, M. Gheorghe, D. Baranašić, W. Santana-Garcia, G. Tan, J. Chèneby, B. Ballester, F. Parcy, A. Sandelin, B. Lenhard, W. W. Wasserman, A. Mathelier, JASPAR 2020: update of the open-access database of transcription factor binding profiles. Nucleic Acids Res. 48, D87–D92 (2020).

48. Y. Liu, X. Du, J. Chen, Y. Jin, L. Peng, H. H. X. Wang, M. Luo, L. Chen, Y. Zhao, Neutrophil-to-lymphocyte ratio as an independent risk factor for mortality in hospitalized patients with COVID-19. J. Infect. (2020), doi: 10.1016/j.jinf.2020.04.002.

49. F. A. Lagunas-Rangel, Neutrophil-to-lymphocyte ratio and lymphocyte-to-C-reactive protein ratio in patients with severe coronavirus disease 2019 (COVID-19): A meta-analysis. J. Med. Virol. n/a (2020), doi: 10.1002/jmv.25819.

50. C. Qin, L. Zhou, Z. Hu, S. Zhang, S. Yang, Y. Tao, C. Xie, K. Ma, K. Shang, W. Wang, D. S. Tian, Dysregulation of immune response in patients with COVID-19 in Wuhan, China. Clin. Infect. Dis. (2020), doi: 10.1093/cid/ciaa248.

51. Y. Zuo, S. Yalavarthi, H. Shi, K. Gockman, M. Zuo, J. A. Madison, C. N. Blair, A. Weber, B. J. Barnes, M. Egeblad, R. J. Woods, Y. Kanthi, J. S. Knight, Neutrophil extracellular traps in COVID-19. JCI insight (2020), doi: 10.1172/jci.insight.138999.

52. S. Matsuyama, M. Ujike, S. Morikawa, M. Tashiro, F. Taguchi, Protease-mediated enhancement of severe acute respiratory syndrome coronavirus infection. Proc. Natl. Acad. Sci. U. S. A. 102, 12543–12547 (2005).

53. D. Panda, E. Gjinaj, M. Bachu, E. Squire, H. Novatt, K. Ozato, R. L. Rabin, IRF1 maintains optimal constitutive expression of antiviral genes and regulates the early antiviral response. Front. Immunol. 10, 1019 (2019).

54. D. Yamane, H. Feng, E. E. Rivera-Serrano, S. R. Selitsky, A. Hirai-Yuki, A. Das, K. L. McKnight, I. Misumi, L. Hensley, W. Lovell, O. González-López, R. Suzuki, M. Matsuda, H. Nakanishi, T. Ohto-Nakanishi, T. Hishiki, E. Wauthier, T. Oikawa, K. Morita, L. M. Reid, P. Sethupathy, M. Kohara, J. K. Whitmire, S. M. Lemon, Basal expression of interferon regulatory factor 1 drives intrinsic hepatocyte resistance to multiple RNA viruses. Nat. Microbiol. 4, 1096–1104 (2019).

55. J. R. Klune, R. Dhupar, S. Kimura, S. Ueki, J. Cardinal, A. Nakao, G. Nace, J. Evankovich, N. Murase, A. Tsung, D. A. Geller, Interferon regulatory factor-2 is protective against hepatic ischemia-reperfusion injury. Am. J. Physiol. – Gastrointest. Liver Physiol. 303, G666–73 (2012).

56. H. Harada, T. Fujita, M. Miyamoto, Y. Kimura, M. Maruyama, A. Furia, T. Miyata, T. Taniguchi, Structurally similar but functionally distinct factors, IRF-1 and IRF-2, bind to the same regulatory elements of IFN and IFN-inducible genes. Cell. 58, 729–739 (1989).

57. A. Hochhaus, X. H. Yan, A. Willer, R. Hehlmann, M. Y. Gordon, J. M. Goldman, J. V. Melo, Expression of interferon regulatory factor (IRF) genes and response to interferon-α in chronic myeloid leukaemia. Leukemia. 11, 933–939 (1997).

58. G. Ren, K. Cui, Z. Zhang, K. Zhao, Division of labor between IRF1 and IRF2 in regulating different stages of transcriptional activation in cellular antiviral activities. Cell Biosci. 5, 17 (2015).

59. A. R. Krarup, M. Abdel-Mohsen, M. H. Schleimann, L. Vibholm, P. A. Engen, A. Dige, B. Wittig, M. Schmidt, S. J. Green, A. Naqib, A. Keshavarzian, X. Deng, R. Olesen, A. M. Petersen, T. Benfield, L. Østergaard, T. A. Rasmussen, J. Agnholt, J. R. Nyengaard, A. Landay, O. S. Søgaard, S. K. Pillai, M. Tolstrup, P. W. Denton, The TLR9 agonist MGN1703 triggers a potent type i interferon response in the sigmoid colon. Mucosal Immunol. 11, 449–461 (2018).

60. F. Liang, G. Lindgren, A. Lin, E. A. Thompson, S. Ols, J. Röhss, S. John, K. Hassett, O. Yuzhakov, K. Bahl, L. A. Brito, H. Salter, G. Ciaramella, K. Loré, Efficient Targeting and Activation of Antigen-Presenting Cells In Vivo after Modified mRNA Vaccine Administration in Rhesus Macaques. Mol. Ther. 25, 2635–2647 (2017).

61. I. E. Galani, V. Triantafyllia, E. E. Eleminiadou, O. Koltsida, A. Stavropoulos, M. Manioudaki, D. Thanos, S. E. Doyle, S. V. Kotenko, K. Thanopoulou, E. Andreakos, Interferon-λ Mediates Non-redundant Front-Line Antiviral Protection against Influenza Virus Infection without Compromising Host Fitness. Immunity. 46, 875–890.e6 (2017).

62. J. Chen, Y. F. Lau, E. W. Lamirande, C. D. Paddock, J. H. Bartlett, S. R. Zaki, K. Subbarao, Cellular Immune Responses to Severe Acute Respiratory Syndrome Coronavirus (SARS-CoV) Infection in Senescent BALB/c Mice: CD4+ T Cells Are Important in Control of SARS-CoV Infection. J. Virol. 84, 1289–1301 (2010).

63. Y.-T. Yen, F. Liao, C.-H. Hsiao, C.-L. Kao, Y.-C. Chen, B. A. Wu-Hsieh, Modeling the early events of severe acute respiratory syndrome coronavirus infection in vitro. J. Virol. 80, 2684–2693 (2006).

64. J. R. Somoza, J. D. Ho, C. Luong, M. Ghate, P. A. Sprengeler, K. Mortara, W. D. Shrader, D. Sperandio, H. Chan, M. E. McGrath, B. A. Katz, The structure of the extracellular region of human hepsin reveals a serine protease domain and a novel scavenger receptor cysteine-rich (SRCR) domain. Structure. 11, 1123–1131 (2003).

65. V. G. Martínez, S. K. Moestrup, U. Holmskov, J. Mollenhauer, F. Lozano, Pharmacol. Rev., in press, doi: 10.1124/pr.111.004523.

66. E. Böttcher-Friebertshäuser, C. Freuer, F. Sielaff, S. Schmidt, M. Eickmann, J. Uhlendorff, T. Steinmetzer, H.-D. Klenk, W. Garten, Cleavage of influenza virus hemagglutinin by airway proteases TMPRSS2 and HAT differs in subcellular localization and susceptibility to protease inhibitors. J. Virol. 84, 5605–5614 (2010).

67. J. E. Park, K. Li, A. Barlan, A. R. Fehr, S. Perlman, P. B. McCray, T. Gallagher, Proteolytic processing of middle east respiratory syndrome coronavirus spikes expands virus tropism. Proc. Natl. Acad. Sci. U. S. A. 113, 12262–12267 (2016).

68. S. Makris, M. Paulsen, C. Johansson, Type I interferons as regulators of lung inflammation. Front. Immunol. 8, 259 (2017).

69. C. J. Britto, N. Niu, S. Khanal, L. Huleihel, J. D. Herazo-Maya, A. Thompson, M. Sauler, M. D. Slade, L. Sharma, C. S. D. Cruz, N. Kaminski, L. E. Cohn, BPIFA1 regulates lung neutrophil recruitment and interferon signaling during acute inflammation. Am. J. Physiol. – Lung Cell. Mol. Physiol. 316, L321–L333 (2019).

70. L. Andzinski, N. Kasnitz, S. Stahnke, C. F. Wu, M. Gereke, M. Von Köckritz-Blickwede, B. Schilling, S. Brandau, S. Weiss, J. Jablonska, Type i IFNs induce anti-tumor polarization of tumor associated neutrophils in mice and human. Int. J. Cancer. 138, 1982–1993 (2016).

71. T. M. Tumpey, C. F. Basler, P. V Aguilar, H. Zeng, A. Solórzano, D. E. Swayne, N. J. Cox, J. M. Katz, J. K. Taubenberger, P. Palese, A. Garc\’\ia-Sastre, Characterization of the Reconstructed 1918 Spanish Influenza Pandemic Virus. Science (80-.). 310, 77–80 (2005).

72. M. Sahoo, L. del Barrio, M. A. Miller, F. Re, Neutrophil Elastase Causes Tissue Damage That Decreases Host Tolerance to Lung Infection with Burkholderia Species. PLOS Pathog. 10, e1004327 (2014).

73. A. Wang, J. Chiou, O. B. Poirion, J. Buchanan, M. J. Valdez, J. M. Verheyden, X. Hou, M. Guo, J. M. Newsome, P. Kudtarkar, D. A. Faddah, K. Zhang, R. E. Young, J. Barr, R. Misra, H. Huyck, L. Rogers, C. Poole, J. A. Whitsett, G. Pryhuber, Y. Xu, K. J. Gaulton, S. Preissl, X. Sun, N. L. Consortium, bioRxiv, in press, doi: 10.1101/2020.04.12.037580.

74. H. Ohbayashi, Neutrophil elastase inhibitors as treatment for COPD. Expert Opin. Investig. Drugs. 11, 965–980 (2002).

75. T. Németh, M. Sperandio, A. Mócsai, Neutrophils as emerging therapeutic targets. Nat. Rev. Drug Discov. 19, 253–275 (2020).

76. R. Qiu, B. G. Sun, J. Li, X. Liu, L. Sun, Identification and characterization of a cell surface scavenger receptor cysteine-rich protein of Sciaenops ocellatus: bacterial interaction and its dependence on the conserved structural features of the SRCR domain. Fish Shellfish Immunol. 34, 810–818 (2013).

77. C. B. Pereira, M. Bocková, R. F. Santos, A. M. Santos, M. M. de Araújo, L. Oliveira, J. Homola, A. M. Carmo, The scavenger receptor SSc5D physically interacts with bacteria through the SRCR-containing N-terminal domain. Front. Immunol. 7, 416 (2016).

